# Modelling and Interpreting Network Dynamics

**DOI:** 10.1101/124016

**Authors:** Ankit N. Khambhati, Ann E. Sizemore, Richard F. Betzel, Danielle S. Bassett

## Abstract

Recent advances in brain imaging techniques, measurement approaches, and storage capacities have provided an unprecedented supply of high temporal resolution neural data. These data present a remarkable opportunity to gain a mechanistic understanding not just of circuit structure, but also of circuit dynamics, and its role in cognition and disease. Such understanding necessitates a description of the raw observations, and a delineation of computational models and mathematical theories that accurately capture fundamental principles behind the observations. Here we review recent advances in a range of modeling approaches that embrace the temporally-evolving interconnected structure of the brain and summarize that structure in a dynamic graph. We describe recent efforts to model dynamic patterns of connectivity, dynamic patterns of activity, and patterns of activity atop connectivity. In the context of these models, we review important considerations in statistical testing, including parametric and non-parametric approaches. Finally, we offer thoughts on careful and accurate interpretation of dynamic graph architecture, and outline important future directions for method development.

---

The increasing availability of human neuroimaging data acquired at high temporal resolution has spurred efforts to model and interpret these data in a manner that provides insights into circuit dynamics [1]. Such data span many distinct imaging modalities and capture inherently different indicators of underlying neural activity, neurotransmitter function, and excitatory/inhibitory balance [2, 3]. Particularly amenable to whole-brain acquisitions, the development of multiband imaging has provided an order of magnitude increase in the temporal resolution of one of the slowest imaging measurements: functional magnetic resonance imaging (fMRI) [4]. Over limited areas of cortex, intracranial electrocorticography (ECoG) complements magnetoencephalography (MEG) and electroencephalography (EEG) by providing sampling frequencies of approximately 2 kHz and direct measurements of synchronized postsynaptic potentials at the exposed cortical surface [5]. In each case, data can be sampled from many brain areas over hours (fMRI) to weeks (ECoG), providing increasingly rich neurophysiology for models of brain dynamics [6], both to explain observations in a single modality, and to link observations across modalities [7].

A common guiding principle across many of these modeling endeavors is that the brain is an interconnected complex system (Fig. 1), and that understanding neural function may therefore require theoretical tools and computational methods that embrace that interconnected structure [8]. The language of networks and graphs has proven particularly useful in describing interconnected structures throughout the world in which we live [9]: from vasculature [10] and genetics [11] to social groups [12] and physical materials [13]. Historically, the application of network science to each of these domains tends to begin with a careful description of the network architecture present in the system, including comparisons to appropriate statistical null models [14]. Descriptive statistics then give way to generative models that support prediction and classification, and eventually efforts focus on fundamental theories of network development, growth, and function [15]. In the context of neural systems, these tools are fairly nascent – with the majority of efforts focusing on description [16], a few efforts beginning to tackle generation and prediction [17–19], and still little truly tackling theory [20, 21].

**FIG. 1.**
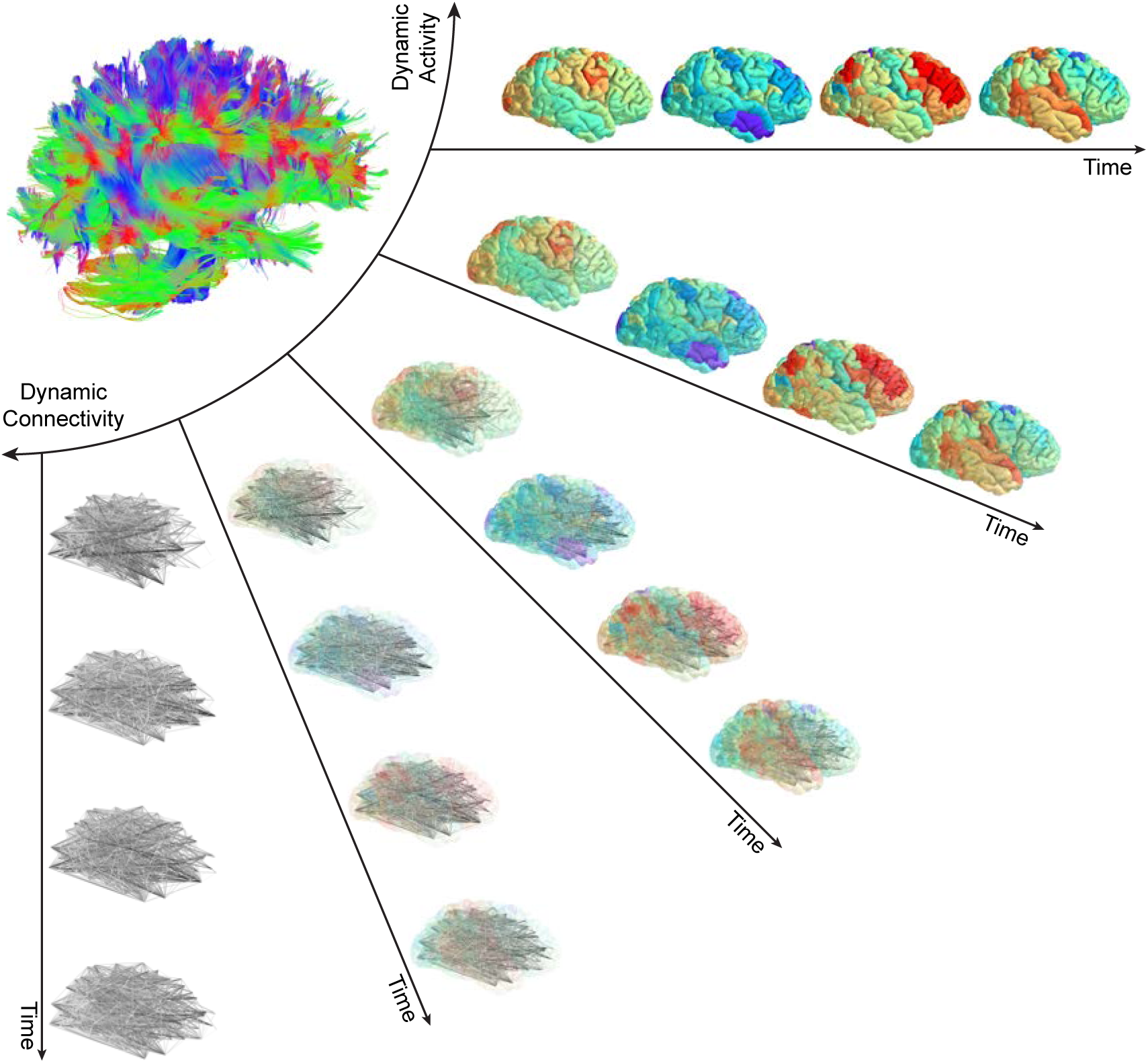
Continuum of functional activity and network dynamics. The structural connectome supports a diverse repertoire of functional brain dynamics, ranging from the patterns of activity across individual brain regions to the patterns of connections interconnecting brain regions. Traditional efforts to understand information processing in the brain have focused on tracking either functional connectivity or functional activity, separately. Novel tools grounded in graph theory, machine learning, signal processing, and algebraic topology now help neuroscientists tease apart the dynamics of functional connectivity and functional activity, concurrently, to begin understanding their complimentary role in neural processing.

The recent extension of network models to the time domain supports the movement from description to prediction (and eventually theory), and also capitalizes on the increasing availability of high-resolution neuroimaging data. Variously referred to as dynamic graphs [22], temporal networks [23], or dynamic networks [24], graph-based models of time-evolving interconnection patterns are particularly well-poised to enhance our understanding of dynamic neural processes from cognition and psychosis [25, 26], to development and aging [27, 28]. Here we review several of these recently developed methods that have been built upon the mathematical foundations of graph theory [29, 30], and we complement a description of the approaches with a discussion of statistical testing and interpretation. To ensure that the treatise is conceptually accessible and manageable in terms of length, we place a particular focus on approaches quantifying meso-scale network architecture; new efforts in local and global-scale network architecture are covered in [31]. Readers are directed elsewhere for recent reviews on multi-scale network models [32], multi-scale biophysical models [33], ICA-based approaches [34], dynamic causal modeling [35], and whole-brain dynamical systems models [6] including the Virtual Brain [36].

The remainder of this review is organized as follows. We begin with a simple description of network (or graph) models that can be derived from functional signals across different imaging modalities. We then describe recent efforts to model network dynamics by considering (i) patterns of connectivity, (ii) levels of activity, and (iii) activity atop connectivity. Next, we discuss statistical testing of network dynamics building on graph null models, time series null models, and other related inferences approaches. We conclude with a discussion of important considerations when interpreting network dynamics, and outline a few future directions that we find particularly exciting.

## GRAPH MODELS OF FUNCTIONAL SIGNALS

Before describing the recent methodological advances in dynamic graph models, and their application to neuroimaging data, it is important to clarify a few definitions. First, we use the term *graph* in the mathematical sense to indicate a graph *G* = (*V, E*) composed of a node set *V* with size *N* and an edge set *E*, and we store this information in an adjacency matrix **A**, whose elements *A_ij_* indicate the strength of edges between nodes [29, 30]. Second, we use the term *model* to indicate a simplified representation of raw observations; a graph model parsimoniously encodes the relationships between system components [37]. Given these two first definitions, it is natural that we use the term *dynamic graph model* to indicate a time-ordered set of graph models of neural data: a single adjacency matrix encodes the pattern of connectivity at a single time point or in a single time window t of data, and the set of adjacency matrices extends that encoding over many time points or many time windows. Dynamic graph models have the disadvantage of ignoring non-relational aspects of the data, but have unique advantages in providing access to a host of computational tools and conceptual frameworks developed by the applied mathematics community over the last few decades. Moreover, dynamic graph models are a singular representation that can be flexibly applied across spatial and temporal scales, thereby supporting multimodal investigations [38, 39] and cross-species analyses [40, 41].

Dynamic graph models can be built from fMRI, EEG, MEG, ECoG, and other imaging modalities using similar principles. First, vertices of the graph (or nodes of the network) need to be chosen, followed by a measure quantifying the strength of edges linking two vertices. In fMRI, nodes are commonly chosen as contiguous volumes either defined by functional or anatomical boundaries [42, 43]. Edge weights are commonly defined by a Pearson correlation coefficient [44]; however, a growing number of studies uses a magnitude squared coherence, to increase robustness to artifacts and to ensure that regional variability in the hemodynamic response function does not create artifactual structure as it can in a correlation matrix [45–47]. In EEG and MEG data, nodes usually represent either sensors or sources obtained after applying source-localization techniques; edges usually represent spectral coherence [48], mutual information [49], phase lag index [50], or synchronization likelihood [51]. In ECoG data, an increasingly popular method to define functional relationships between sensors is a multi-taper coherence [52, 53]. Note: While neuron-level recordings are not the focus of this exposition, the tools we describe here are equally applicable to dynamic graphs in which neurons are represented as nodes [54], and in which relationships between neurons are summarized in, for example, shuffle-corrected cross-correlograms [55–58].

After nodes have been chosen and the edge measure defined, the widely-accepted approach for generating a dynamic graph model is to delineate time windows, where the pattern of functional connectivity in each time window is encoded in an adjacency matrix. Choosing the size of the time window is important. Short windows may hamper accurate estimates of functional connectivity within the frequency band of interest [59, 60], while long windows may only reflect the time-invariant network structure of the data [61–63]. Intuitively, to achieve accurate estimates of covariation in slow fluctuations, one would like to include several cycles of the signal [59, 64]. In our recent work, we demonstrated that short time windows may better reflect individual differences while long time windows may reflect network architectures that are reproducible over iterative measurement [60]. We also suggest that a reasonable method for choosing a time window of interest is to maximize the variability in the network’s flexible reconfigurations over time (see later sections for further details). Prior work has used similar approaches to suggest optimal time windows on the order of a few 10’s of seconds for human BOLD data, and 1 s for ECoG data [52, 53, 60, 65].

## MODELING NETWORK DYNAMICS

The construction process described in the previous section provides a dynamic graph model from which one can begin to infer organizational principles and their temporal variation. In this section, we describe methods that build on these models to characterize time-evolving patterns of connectivity. We then describe a set of related methods that characterize time-evolving patterns of activity, and we conclude this section by describing methods that explicitly characterize how activity occurs atop connectivity.

### Considering patterns of connectivity

Time-varying graph dynamics can be thought of as a type of system evolution (Fig. 2). When considering canonical forms of evolution, one quite naturally thinks about modularity [66]: the nearly decomposable nature of many adaptive systems that supports their potential for evolution and development [67–69]. Modularity is a consistently observed characteristic of graph models of brain function [70, 71], where it is thought to facilitate segregation of function [72], learning without forgetting [73], and potential for rehabilitation after injury [74].

**FIG. 2.**
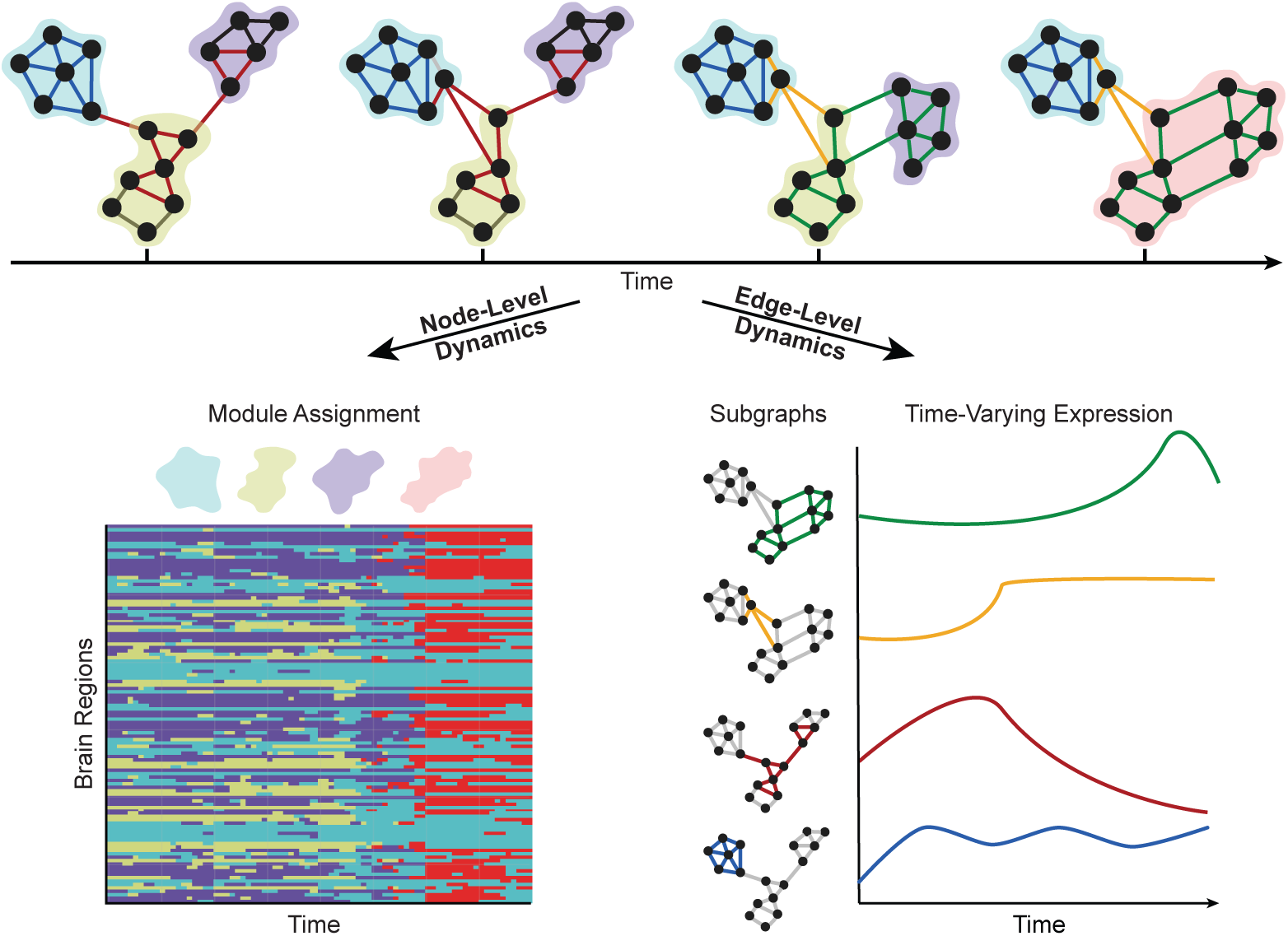
Dynamic network modules and subgraphs. (*Top*) Network science enables investigators to study dynamic architecture of complex brain networks in terms of the collective organization of nodes and of edges. Clusters of strongly interconnected nodes are known as modules, and clusters of co-varying edges are known as subgraphs. Nodes/edges of the same module/subgraph are shaded by color. Each module represents a functionally cohesive collection of nodes, and each node may only be a member of a single module. Each subgraph is a recurring pattern of edges that link information between nodes at the same points in time, and each edge can belong to multiple subgraphs. (*Bottom Left*) Dynamic community detection assigns nodes to time-varying modules. Nodes may shift their participation between modules over time based on the demands of the system. (*Bottom Right*) Non-negative matrix factorization pursues a parts-based decomposition of the dynamic network into subgraphs and time-varying coefficients, which quantify the level of expression of each subgraph over time.

The ability to assess time-varying modular architecture in brain graphs is critical for an understanding of the exact trajectories of network reconfiguration that accompany healthy cognitive function and development [75], as well as the identification of altered trajectories characteristic of disease [76]. Yet, there are several computational challenges that must be addressed. The simplest method to assess time-varying modular architecture is to identify modules in each time window, and then develop statistics to characterize their changes. However, identifying changes in modules requires that we have a map from a module in one time window to itself in the next time window. Such a map is not a natural byproduct of methods applied to individual time windows separately; due to the heuristic nature of the common community detection algorithms [77–79], a module assigned the label of *module 1* in time window *l* need not be the same as the module assigned the same label in time window *r*. The historic Hungarian algorithm (developed in 1955) can be used to attempt a re-labeling to create an accurate mapping [80, 81], but the algorithm fails when ties occur, and is not parameterized to assess mappings sensitive to module-to-module similarities occurring over different time scales.

#### Dynamic Community Detection

A recent solution to these problems lies in transforming the ordered set of adjacency matrices that compose a dynamic graph model into a multilayer network [82]. Here, the graph in one time window is linked to the graph in adjacent time windows by identity edges that connect a node in one time window to itself in neighboring time windows [83, 84]. Then, one can identify modules – and their temporal variation – by maximizing a multilayer modularity quality function:

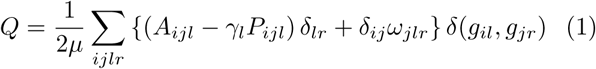
where *A_ijl_* is the weight of the edge between nodes *i* and *j* in a time window *l*; the community assignment of node *i* in layer *l* is *g_il_*, the community assignment of node *j* in layer *r* is *g_jr_*, and *δ*(*g_il_, g_jr_*) = 1 if *g_il_* = *g_jr_* and 0 otherwise; the total edge weight is 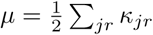, where *κ_jl_* = *k_jl_* + *c_jl_* is the strength of node *j* in layer *l, k_jl_* is the intra-layer strength of node *j* in layer *l*, and 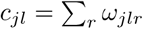 is the inter-layer strength of node *j* in layer *l*. The variable *P_ijl_* is the corresponding element of a specified null model, which can be tuned to account for different characteristics of the system [13, 85, 86]. The parameter *γ_l_* is a structural resolution parameter of layer *l* that can be used to tune the number of communities identified, with lower values providing sensitivity to large-scale community structure and higher values providing sensitivity to small-scale community structure. The strength of the identity link between node *j* in layer *r* and node *j* in layer *l* is *ω_jlr_*, which can be used to tune the temporal resolution of the identified module reconfiguration process, with lower values providing sensitivity to high-frequency changes in community structure and higher values providing sensitivity to low-frequency changes in community structure [87].

The dynamic community detection approach has several strengths. First, it solves the matching problem that defines which module in one time window “is the same as” which module in another time window. Second, it provides tuning parameters that enable one to access information about both fine and coarse topological scales of network reconfiguration (*γ*), as well as both fine and coarse temporal scales of network reconfiguration (*ω*). Third, the formulation allows one to construct and incorporate hypothesis-specific null models (**P**). Fourth, unlike statistically-driven methods based on principle components analysis or independent components analysis, there are no requirements for modules to be completely independent from one another. Fifth, the method provides natural ways to assess overlapping community structure, either by examining the probability that nodes are assigned to a given community over multiple optimizations of the modulatiy quality function, or by extending the tool to identify communities of edges [88–90]. Sixth, generative models for module reconfiguration processes are beginning to facilitate the potential transition from description to prediction and theory [91, 92]. These advantages have proven critical for studies of network dynamics accompanying working memory [24], attention [93], mood [94], motor learning [95, 96], reinforcement learning [97], language processing [98, 99], inter-task differences [60, 65], normative development and aging [100, 101], inter-frequency relationships [102], and behavioral chunking [86].

#### Non-Negative Matrix Factorization

Of course, the clustering of nodes into functionally-cohesive modules is just one of potentially many organizational principles characterizing dynamic brain networks. Dynamic community detection provides a lens on the dynamics of node-level organization in the network, but it remains agnostic to the dynamics of edges that link nodes within and between modules. Recent advances in graph theoretic tools based on machine learning can provide insights into additional constraints on the evolution of brain systems [28, 103, 104] by addressing open questions such as: How are the edges linking network nodes changing with time? Do all edges reorganize as a cohesive group, or are there smaller clusters of edges that reorganize at different rates or in different ways? Could the same edge link two nodes of the same module at one point in time and link two nodes of different modules at another point in time?

One set of tools that can begin answering these questions is an unsupervised machine learning approach known as non-negative matrix factorization (NMF) [105]. NMF has previously been applied to neuroimaging data to extract structure in morphometric variables [106], tumor heterogeneity [107], and resting state fMRI [108]. In the context of dynamic graph models, NMF objectively identifies clusters of co-evolving edges, known as *subgraphs*, in large, temporally-resolved data sets [109]. Conceptually, subgraphs are mathematical basis functions of the dynamic brain graph whose weighted linear combination – given by a set of time-varying basis weights or expression coefficients for each subgraph – reconstructs a repertoire of graph configurations observed over time. In contrast to the hard-partitioning of nodes into discrete modules in dynamic community detection, NMF pursues a soft-partitioning of the network such that all graph nodes and edges participate to varying degree in each subgraph – which is represented by a weighted adjacency matrix (see [109] for in-depth comparison of network modules and network subgraphs).

To apply NMF to the dynamic graph model, the edges in the *N* × *N* × *T* dynamic adjacency matrix must be non-negative and unraveled into a *N*(*N* − 1)/2 × *T* network configuration matrix **Â**. Next, one can minimize the *L*2-norm reconstruction error between **Â** and the matrix product of two non-negative matrices **W** – an *N*(*N* − 1)/2 × *m* subgraph matrix – and **H** – an *m* × *T* time-varying expression coefficients matrix – such that:

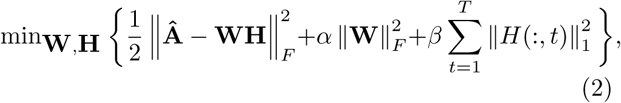
where ║·║*_F_* is the Frobenius norm operator, ║·║_1_ is the L1 norm operator, *m* ∈ [2, min(*N*(*N* − 1)/2, *T*) − 1] is a rank parameter of the factored matrices that can be used to tune the number of subgraphs to identify, *β* is a tunable penalty weight to impose sparse temporal expression coefficients, and *α* is a tunable regularization of the edge strengths for subgraphs [110]. To tune these parameters without overfitting the model to dynamic network data, recent studies have employed parameter grid search [28] and random sampling approaches [104, 109].

NMF has distinct advantages over other unsupervised matrix decomposition approaches such as PCA [111]. First, the non-negativity constraint in the NMF approach means that subgraphs can be interpreted as additive components of the original dynamic network. That is, the relative expression of different subgraphs can be judged purely based on the positivity of the expression coefficients during a given time window. Second, NMF does not make any explicit assumptions about the orthogonality or independence of the resulting subgraphs, which provides added flexibility in uncovering network components with overlapping sub-structures that are specific to different brain processes. Recent applications of NMF to characterize network dynamics at the edge-level have led to important insights into the evolution of executive networks during healthy neurodevelopment [28] and have uncovered putative network components of function and dysfunction in medically refractory epilepsy [104].

### Considering levels of activity

The approaches described in the previous section seek to characterize mesoscale structure in dynamic graph models in which nodes represent brain areas and edges represent functional connections. Yet, one can construct alternative graph models from neuroimaging data to test different sorts of hypotheses. In particular, a set of new approaches have begun to be developed for understanding how brain states evolve over time: where a state is defined as a pattern of activity over all brain regions [112–116], rather than as a pattern of connectivity. This notion of brain state is one that has its roots in the analysis of EEG and MEG data, where the voltage patterns across a set of sensors or sources has been referred to as a microstate [117]. The composition and dynamics of microstates predict working memory performance [118] and are altered in disease [119]. Emerging graph theoretical tools have become available to study how such states evolve into one another. These activity-centric approaches are reminiscent of multi-voxel pattern analysis (MVPA) approaches in the sense that the object of interest is a multi-region pattern of activity [120, 121]; yet, they differ from MVPA in that they explicitly use computational tools from graph theory to understand complex patterns of relationships between states.

#### Time-by-Time Graphs

Perhaps the simplest example of such an approach is the construction of so-called *time-by-time* networks [122–124]: a graph whose nodes represent moments in time, and whose edges represent similarities between moment pairs (Fig. 3A-B). For example, in the context of BOLD fMRI, a node could represent a repetition time sample (TR), and an edge between two TRs could indicate a degree of similarity or distance between the brain state at TR *l* and the brain state at TR *r*. We will represent this graph as the adjacency matrix **T** which is of dimension *T* × *T*, in contrast to the traditionally studied adjacency matrix **A** which is of dimension *N* × *N*.

**FIG. 3.**
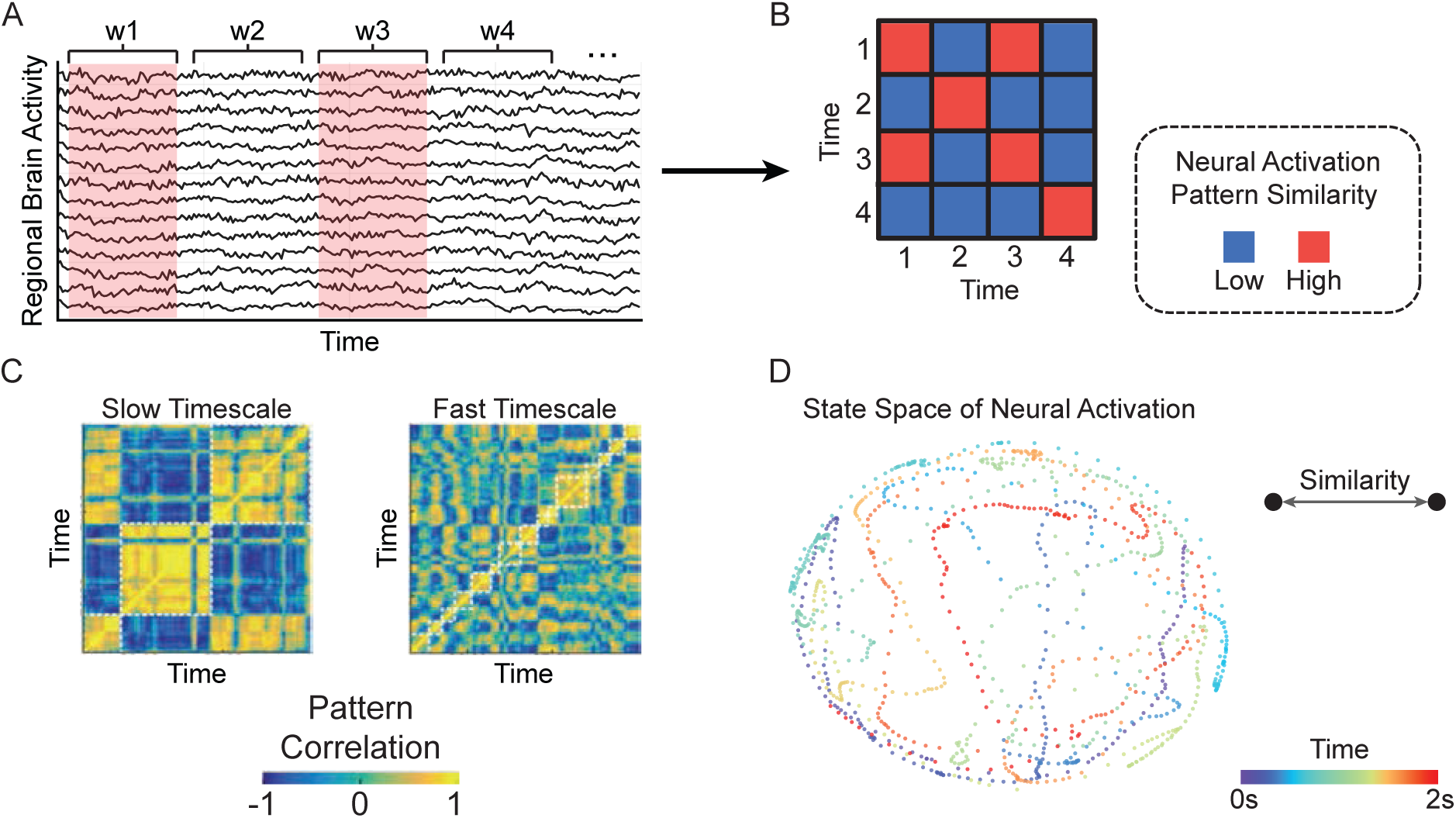
State space of brain activity patterns. (*A*) The time-by-time graph captures similarities in neural activation patterns between different points in time. In practice, one can compute the average brain activity for individual brain regions within discrete time windows and compare the resulting pattern of activation between two time windows using a similarity function, such as the Pearson correlation. (*B*) The resulting time-by-time graph has an adjacency matrix representation in which each time window is a node and the neural activation similarity between a pair of time windows is an edge. (*C*) Clustering tools based on graph theory or machine learning can identify groups of time windows – or states – with similar patterns of neural activation. By parametrically varying the number of clusters, or their size, one can examine the dynamic states over multiple time scales (Adapted from [126]). (*D*) Example two-dimensional projection of a time-by-time graph derived from ECoG during a 2 s, baseline interval. Each node represents a 2 ms time window and is shaded based on its occurrence in the 2 s interval. The spatial proximity between pairs of nodes represents the similarity in their neural activation pattern. The depicted state space demonstrates an interleaved trajectory during the brief baseline period.

After constructing a time-by-time network, one can apply graph theoretical techniques to extract the community structure of the graph, to identify canonical states, and to quantify the transitions between them (Fig. 3C-D). Efforts in this vein have identified different canonical states in rest [122, 123] versus task [124], and observed that flexible transitions between states change over development [122] and predict individual differences in learning [124]. While these studies have focused on the cluster structure of time-by-time graphs, other metrics – including local clustering and global efficiency – could also be applied to these networks to better understand how the brain traverses states over time. In related work, reproducible temporal sequences of states have been referred to as lag threads [125], and boundaries between states have offered important insights into the storage and retrieval of events in long-term memory [126].

#### Topological Data Analysis: Mapper

A conceptually similar approach begins with the same underlying data type (an *N* × *T* matrix representing regional activity magnitudes as a function of time) and applies tools from algebraic topology to uncover meaningful – and statistically unexpected structure – in evolving patterns of neural activity [127].

Generally, these raw data arise from sampling a possibly high-dimensional manifold which describes all possible occurrences of the observed data type – implying that the global shape or topological features of this manifold could inform our understanding of processes specific to the system. In mathematics, multiple methods exist for returning topological information about a manifold. One such method is to construct a simplified object from our topological space *X*, called the Reeb graph [128], which captures the evolution of the connected components within level sets of a continuous function *f*: *X* → ℝ. As an example, if our topological space *X* is a torus (Fig. 4A), then we can use the height function *h*: *X* → ℝ, so the level sets, *h*^−1^(*c*) for *c* ∈ ℝ, are horizontal slices of *X* at height *c*. Note each horizontal slice has one or two (or zero) connected components. As we move from one slice to the next, we record only the critical points at which the number of connected components *changes*, and how the connected components split or merge at these points. With nodes as critical points and edges indicating connected component evolution, the Reeb graph succinctly reflects these topological features (Fig. 4A, right).

**FIG. 4.**
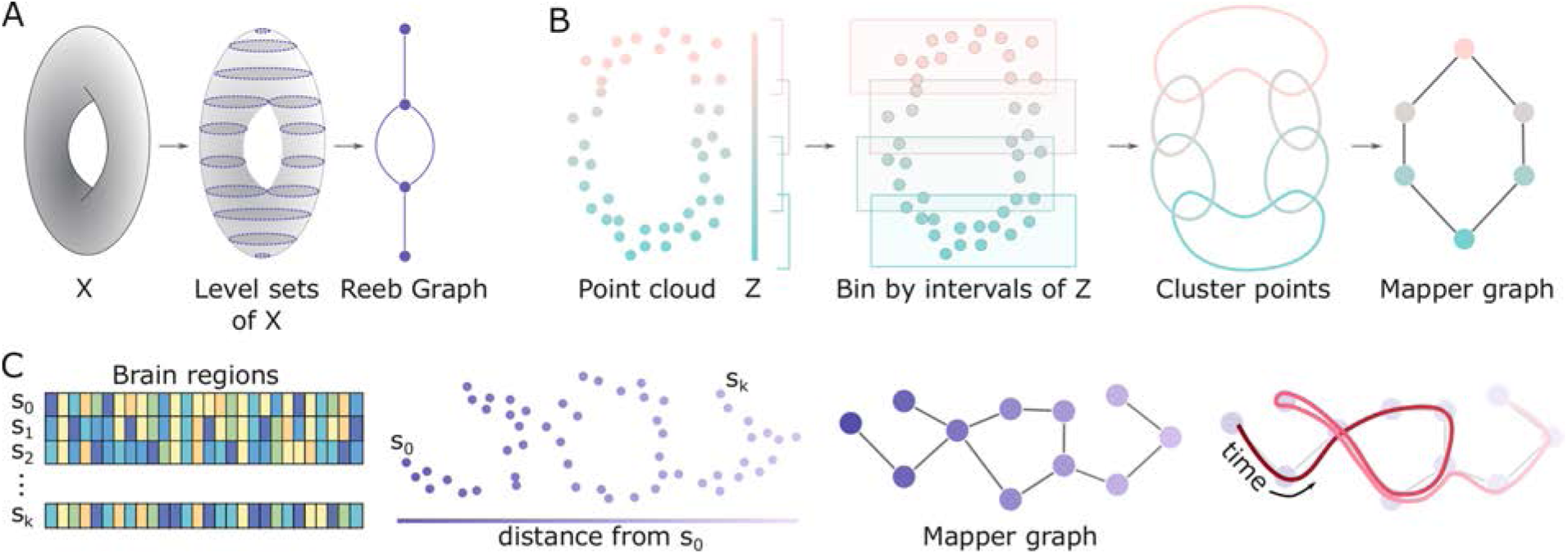
Mapping temporal structure with algebraic topology. (*A*) Schematic of Reeb graph construction. Given a topological space *X*, here a torus, the Reeb graph is constructed by examining the evolution of the level sets from *h*(*X*). (*B*) Illustration of the Mapper algorithm. Beginning with a point cloud and parameter space *Z*, points are binned and clustered. Resulting clusters are collapsed to nodes in the final Mapper graph, and edges between nodes exist if the two corresponding clusters share points from the original point cloud. (*C*) Example use of topological data analysis for dynamic networks. The initial point cloud here is the collection of brain states across time and parameter space the distance from the initial state of the system. Following the path of time (right, red curve) on the Mapper graph may yield insights to system evolution.

While we would prefer a simple summary such as the Reeb graph for neural processes, we record noisy point clouds instead of nice manifolds. One approach for computing summary objects such as this in the presence of noisy data is called Mapper [129], which begins with a point cloud *Y* and two functions: (i) as before, *f* : *Y* → *Z* for some parameter space *Z*, and (ii) a distance metric on *Y* (Fig. 4B, left). We then choose a *cover* of *Z*, or a collection of open sets *U_α_* with *Z* ⊆ **∪***_α_U_α_*. If *Z* is a subset of the real line, in practice we often use a small number of intervals with fixed length. We look then to the collection of points of *Y* falling within *f*^−1^(*U_α_*) for each *α*. In other words, we bin points in *Y* based on their associated *z* ∈ *Z*, and cluster these points using the chosen distance metric. This is analogous to taking level sets and determining connected components when constructing the Reeb graph. The last step creates the output graph (here we call the Mapper graph) – defined using a single node for each cluster and laying edges between nodes whose corresponding clusters share at least one point *y* ∈ *Y*.

The Mapper algorithm or similar methods have been used to stratify disease states and identify patient subgroups [130–132], conduct proteomics analyses [133, 134], and compare brain morphology [135, 136]. These methods are now ripe for application in dynamic networks, as discussed in this review. One possible avenue to describe the shape of a neural state space might involve modeling the states of brain activity {*s*_0_, *s*_1_, …, *s_k_*} as points in a cloud and defining the association parameter *Z* as the Euclidean distance between the initial state, *s*_0_, and all other states (Fig. 4C, left). By binning and clustering points in the cloud, we could recover a graph summarizing the topological features of the state space traversed. This approach would enable us to track whether the system re-visits previously encountered states or enters novel states, as evidenced by loops, dead zones, and branches of the traversed path itself (Fig. 4C, right). Additional possibilities include using density or eccentricity as parameters, or combining these which would yield a higher dimensional output [129].

### Considering activity atop connectivity

The approaches described thus far address either evolving patterns of connectivity (dynamic community detection and non-negative matrix factorization) or evolving patterns of activity (time-by-time networks and Reeb graphs). A natural next question is whether these two perspectives on brain function can be combined in a way that provides insights into how activity occurs atop connectivity. In this section we will describe two such recently-developed approaches stemming respectively from applied mathematics and engineering – annotated graphs and graph signal processing – that have been recently applied to multimodal neuroimaging data to better understand brain network dynamics.

#### Annotated graphs

The traditional composition of a graph includes identical nodes and non-identical (weighted) edges, and this composition is therefore naturally encoded in an adjacency matrix **A**. What is not traditionally included in graph construction – and also not naturally encoded in an adjacency matrix – is any identity or weight associated with a node. Yet, in many systems, nodes differ by size, location, and importance in a way that need not be identically related to their role in the network topology. Indeed, understanding how a node’s features may help to explain its connectivity, or how a node’s connectivity may constrain its features can be critical for explaining a system’s observed dynamics and function.

Annotated graphs are graphs that allow for scalar values or categories (*annotations*) to be associated with each node (Fig. 5A). These graphs are represented by both an adjacency matrix **A** of dimension *N* × *N*, and a vector *x* of dimension *N* × 1. Characterizing annotated graph structure, and performing statistical inference, requires an expansion of the common graph analysis toolkit [137, 138]. In an unannotated graph, network communities are composed of densely interconnected nodes. In an annotated graph, network communities are composed of nodes that are both densely interconnected *and* have similar annotations. Recent efforts have formalized the study of community structure in such graphs by writing down the probability of observing a given annotated graph as a product of the probability of observing that exemplar of a weighted stochastic block model and the probability of observing the annotations [139]. That is, assuming independence between the annotation *x* and the graph **A** with block structure *θ* and community partition *z*:

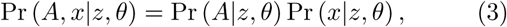
where the first term of the right side of the equation accounts for the probability of observing the graph given the community structure under the assumptions of the weighted stochastic block model. This term relies on the assumption that interconnected nodes are likely to be in the same community. The second term accounts for the probability of observing the continuously valued annotations given the community structure [140]. This term relies on the assumption that nodes with similar annotations are more likely to be in the same community. Methods are available to fit this model to existing data to extract community structure and quantify the degree to which that community structure aligns with node-level annotations. These methods function by finding the community partition *z* that maximizes the probability of Eq. 3.

**FIG. 5.**
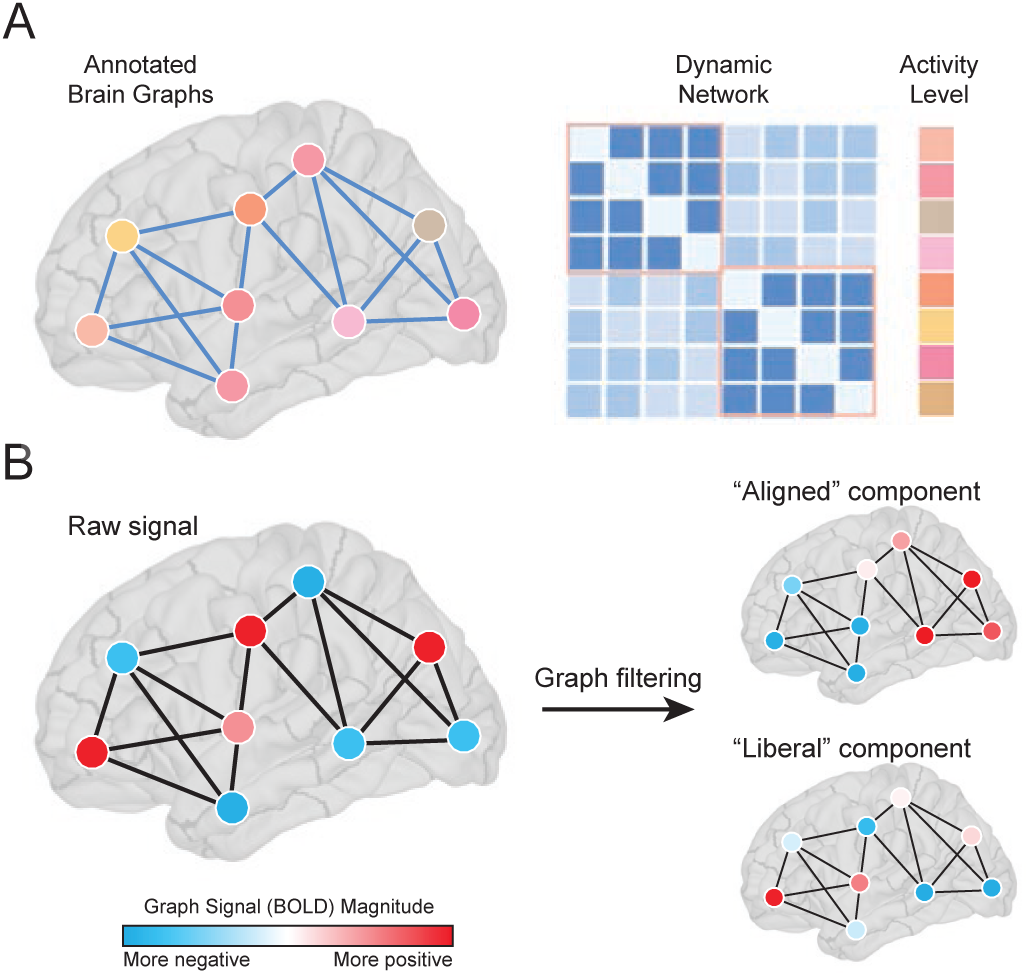
Brain activity on brain graphs. (*A*) Annotated graphs enable the investigator to model scalar or categorical values associated with each node. These graphs are represented by both an adjacency matrix **A** of dimension *N* × *N*, and a vector *x* of dimension *N* × 1 (Adapted from [140]). (*B*) Graph signal processing allows one to interpret and manipulate signals atop nodes in a mathematical space defined by their underlying graphical structure. A graph signal is defined on each vertex in a graph. For example, the signal could represent the level of BOLD activity at brain regions interconnected by a network of fiber tracts. Graph filtering decomposes that signal into components aligned/misaligned with the eigenvectors of the network. A signal is aligned if directly connected vertices have similar amplitude/sign and misaligned (or “liberal”) when the opposite is true

Initial efforts applying these tools to neuroimaging data have demonstrated that discrepancies between BOLD magnitudes and the community structure of functional connectivity patterns predicts individual differences in the learning of a new motor skill [140]. Future efforts could build on these preliminary findings to incorporate different types of annotations, such as measurements of time series complexity [141], gray matter density, cortical thickness, oxidative metabolism [142], gene expression [143], or cytoarchitectural characteristics [144], to better understand the relationships between these regional characteristics and inter-regional estimates of structural or functional connectivity.

#### Graph Signal Processing

A discrete-time signal consists of a series of observations, and can be operated on and transformed using tools from classical signal processing, e.g. filtered, denoised, downsampled, etc. [145]. Many signals, however, are defined on the vertices of a graphs and therefore exhibit interdependencies that are contingent upon the graph’s topological organization. Graph signal processing (GSP) is a set of mathematical tools that implement operations from classical signal processing while simultaneously incorporating and respecting the graphical structure underlying the signal [146] (Fig. 5B). While GSP, in general, has been widely applied for purposes of image compression [147] and semisupervised learning [148] (among others), only recently has it been used to investigate patterns in neuroimaging data [149–151].

One particularly interesting application involves using graph Fourier analysis to study the relationship of a graph signal to an underlying network. This approach, analogous to classical Fourier analysis, decomposes a (graph) signal along a set of components, each of which represents a different mode of spatial variation (graph frequency) with respect to the graph’s toplogical structure. These modes are given by an eigendecomposition of the graph Laplacian matrix, **L** = **D** − **A**, where **D** = diag(*s*_1_, …, *s_N_*) and where *s_i_* = Σ*_j_ A_ij_*. The eigendecomposition results in a set of ordered eigenvalues, λ_0_ ≤ λ_1_ ≤ … ≤ λ_*N*−1_, and corresponding eigenvectors, **Λ** = [Λ_0_, …, Λ_*N*−1_]. The “alignment” of each eigenvector with respect to the underlying network is positive or negative depending upon whether regions with the same or opposite sign are directly connected. Note that the eigendecomposition can also be carried out on the original connectivity matrix, **A**, as well, which might improve interpretation in some cases.

One recent study applied graph Fourier analysis to neuroimaging data to study the relationship of regional activity (BOLD) and inter-regional white-matter networks [152]. In this study, the authors designed a group of graph filters from the eigenvectors most/least aligned with the connectivity matrix, **A**, which decomposed the vector time series of BOLD activity, **B** = [b_1_(*t*), …, b_*N*_(*t*)], into filtered time series 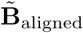 and 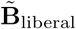, representing those components of the unfiltered signal that were aligned and misaligned with **A**. Interestingly, the extent to which the most ”liberal” (misaligned) signals across the brain aligned with the anatomical networks was associated with cognitive switching costs during a visual perception task, potentially indicating that brain signals which conform to individual anatomy promotes cognitive flexibility.

These recent applications of graph signal processing [149, 151, 152] highlight its utility for studying neural systems at the network level. Nonetheless, there are open methodological and neurobiological questions. For instance, the graph filtering procedure described above operates on the BOLD activity at each instant, independent of the activity at all other time points, and also assumes that the underlying network structure is fixed (i.e. static). Extending the framework to explicitly incorporate the dynamic nature of the signal/network remains an unresolved issue [153]. Also, while graph signal processing operations can identify network-level correlates of behavioral relevance, how these operators are realized neurobiologically is also unclear. Future work could be directed to investigate these and other open questions.

## STATISTICAL TESTING OF NETWORK DYNAMICS

When constructing and characterizing dynamic graph models and other graph-based representations of neuroimaging data, it is important to determine whether the dynamics that are observed are expected or unexpected in an appropriate null model. While no single null model is appropriate for every scientific question, there are a family of null models that have proven particularly useful in testing the significance of different features of brain network dynamics. Generally speaking, these null models fall into two broad categories: those that directly alter the structure of the graph, and those that alter the structure of the time series used to construct the graph. In this section, we will describe common examples of both of these types of null models, and we will also discuss statistical approaches for identifying data-driven boundaries between time windows used to construct the graphs.

### Graph null models

The construction of graph-based null models for statistical inference has a long history dating back to the foundations of graph theory [29, 30]. Common null models used to query the architecture of static graphs include the Erdos-Renyi random graph model and the regular lattice [154–156] – two benchmark models that have proven particularly useful in estimating small-worldness [157, 158]. To address questions regarding network development and associated physical constraints, both spatial null models [85, 159] and growing null models [18] have proven particularly useful. In each case, the null model purposefully maintains some features of interest, while destroying others.

When moving from static graph models to dynamic graph models, one can either devise model-based nulls or permutation-based nulls. Because generative models of network reconfiguration are relatively new [91, 92], and none have been validated as accurate fits to neuroimaging data, the majority of nulls exercised in dynamic graph analysis are permutation-based nulls. In prior literature, there are three features of a dynamic graph model that are fairly straightforward to permute: the temporal order of the adjacency matrices, the pattern of connectivity within any given adjacency matrix, and (for multilayer graphs) the rules for placing and weighting the inter-layer identity links [95]. Permuting the order of time windows uniformly at random in a dynamic graph model is commonly referred to as a *temporal null model*. Permuting the connectivity within a single time window (that is, permuting the location of edges uniformly at random throughout the adjacency matrix) is commonly referred to as a *connectional null model*. For multilayer networks in which the graph in time window *t* is linked to the graph in time window *t* + 1 and also to the graph in time window *t* − 1, one can permute the identity links connecting a node with itself in neighboring time windows uniformly at random. This is commonly referred to as a *nodal null model*.

These permutation-based null models are important benchmarks against which to compare dynamic graph architectures because they separately perturb distinct dimensions of the dynamic graph’s structure. The temporal null model can be used to test hypotheses regarding the nature of the temporal evolution of the graph; the connectional null model can be used to test hypotheses regarding the nature of the intra-window pattern of functional connectivity; and the nodal null model can be used to test hypotheses regarding the importance of regional identity in the observed dynamics [87]. It will be interesting in the future to expand this set of non-parametric null models to include parametric null models that have been carefully constructed to fit generalized statistical structure of neuroimaging data.

### Time series null models

Dynamic graph null models are critical for practitioners conversant in graph theory to understand the driving influences in their data. Yet, others trained in dynamical system theory may wish to understand better how characteristics of the time series drive the observed time evolving patterns of functional connectivity that constitute the dynamic graph model [160]. For these questions, surrogate data time series are particularly useful.

Two historically relevant surrogate data generation techniques for time series are the Fourier transform (FT) surrogate and the amplitude adjusted Fourier transform (AAFT) surrogate. Both methods preserve the mean, variance, and autocorrelation function of the original time series, by scrambling the phase of time series in Fourier space [161]. First, we assume that the linear properties of the time series are specified by the squared amplitudes of the discrete Fourier transform

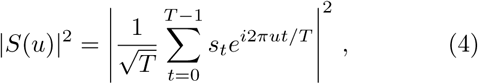
where *s_t_* denotes an element in a time series of length *T* and *S_u_* denotes a complex Fourier coefficient in the Fourier transform of *s*. We can construct the FT surrogate data by multiplying the Fourier transform by phases chosen uniformly at random and transforming back to the time domain:

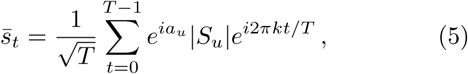
where *a_u_* ∈ [0, 2*π*) are chosen independently and uniformly at random. This approach has proven useful for characterizing brain networks in prior studies [44, 87]. The AAFT extends the FT surrogate by also retaining the amplitude distribution of the original signal [162].

An important feature of these approaches is that they can be used to alter nonlinear relationships between time series while preserving linear relationships (applying an identical shuffling to both time series), or they can be used to alter both linear and nonlinear relationships between time series (applying independent shufflings to each time series) [161]. Other methods that are similar in spirit include those that generate surrogate data using stable vector autoregressive models [168] that approximately preserve the power and cross-spectrum of the actual time series [169]. In each case, after creating surrogate data time series, one can re-apply time window boundaries and extract functional connectivity patterns in each time window to create dynamic graph models. Then one can compare statistics of the dynamic graph models constructed from surrogate data time series with the statistics obtained from the true data [170].

### Change points

Thus far, our discussion regarding statistical inference on dynamic graph models presumes that one has previously determined a set of time windows of the observed time series in which to measure functional connectivity [59, 60]. The common approach is to choose a single time window size, and then to apply it either in a nonoverlapping fashion or in an overlapping fashion [28, 95]. There are several benefits to this approach, including effectively controlling for the influence of the time series length on the observed network architecture. However, one disadvantage of the fixed-window-length approach is that one may be insensitive to changes in the neurophysiological state that occur at non-regular intervals. To address this issue, several methods have been proposed to identify *change points* – points in time where the probability distribution of a time series changes – both in patterns of connectivity [171, 172] and in patterns of activity [173, 174]. These methods employ a data-driven framework to identify irregularly sized time windows before constructing brain graphs using the modeling approaches discussed earlier. Such change point-based network models can potentially enhance tracking of network dynamics alongside changes in cognitive state.

## INTERPRETING NETWORK DYNAMICS

At the conclusion of any study applying the modeling techniques described in this review, it is important to interpret the observed network dynamics within the context in which the data was acquired. When studying network dynamics during a task, assessing the correlation between brain and behavior – both raw behavioral data and parameter values for models fit to the behavioral data – can help the investigator infer network mechanisms associated with task performance [21]. Relevant questions include which network dynamics accompany which sorts of cognitive processes or mental states. For example, recent observations point to a role for frontal-parietal network flexibility in motor learning [96], reinforcement learning [97], working memory [24], and cognitive flexibility [24]. In the case of time-by-time graphs, it is also relevant to link brain states to not only processing functions but also to representation functions, for example by combining local MVPA analyses with global graph analyses. Finally, in both task and rest studies, it is also useful to determine the relationship between brain network dynamics and online measurements of physiology such as pupil diameter [175], galvanic skin response, or fatigue [94].

Outside of the context of a single study, it is important to build an intuition for what types of neurophysiological mechanisms may be driving certain types of brain network reconfiguration – irrespective of whether the subject is performing a task or simply resting inside the scanner. Particularly promising approaches for pinpointing neurophysiological drivers include pharma-fMRI studies, which suggest that distinct neurotransmitters may play important roles in driving network dynamics (Fig. 6). Using an NMDA-receptor antagonist, Braun and colleagues demonstrated that network flexibility – assessed from dynamic community detection – is increased relative to placebo, suggesting a critical role for glutamate in fMRI-derived brain network dynamics [26]. Preliminary data also hint at a role for seratonin by linking a positive effect with network flexibility [94], a role for norepinephrine by linking pupil diameter to network reconfiguration [175], and a role for stress-related corticosteroids and catecholamines in facilitating reallocation of resources between competing attention and executive control networks [176]. However, the relative impact of dopamine, seratonin, glutamate, and norepinephrine on network reconfiguration properties remains elusive, and therefore forms a promising area for further research.

**FIG. 6.**
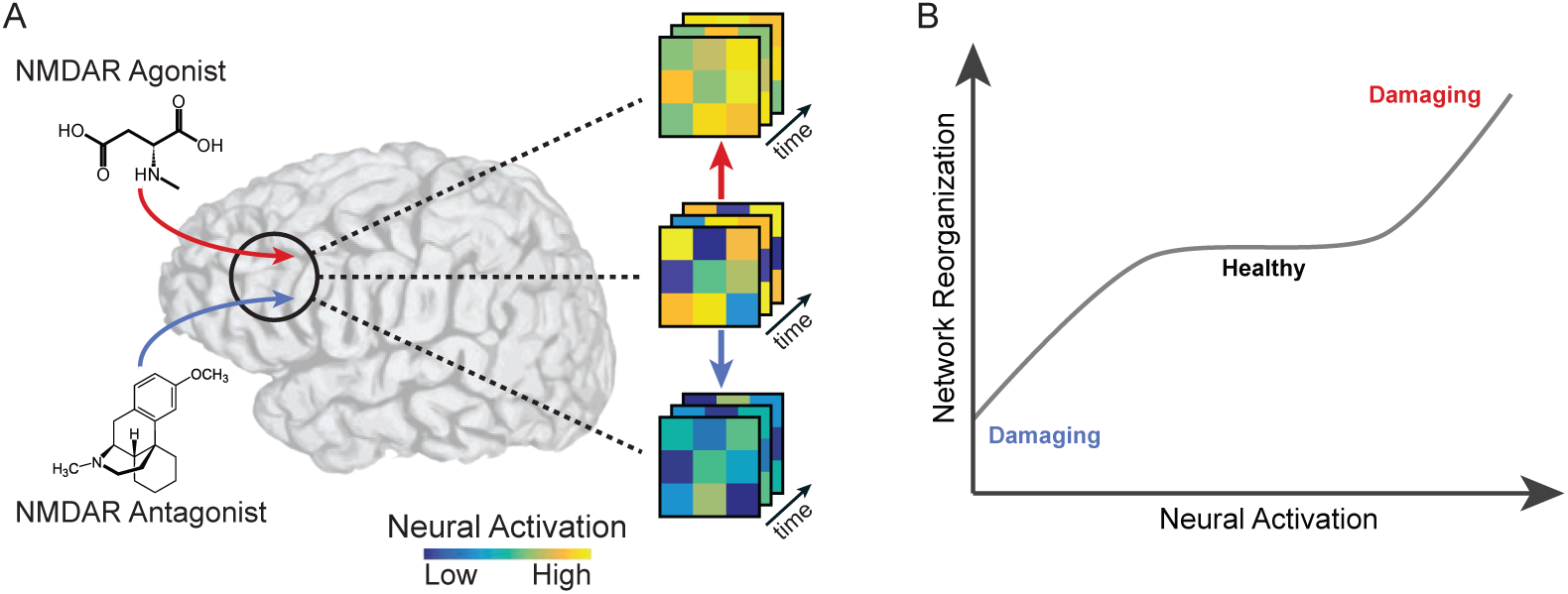
Pharmacologic modulation of network dynamics. (*A*) By blocking or enhancing neurotransmitter release through pharmacologic manipulation, investigators can perturb the dynamics of brain activity. For example, an NMDA receptor agonist might hyper-excite brain activity [163–165], while a NMDA receptor antagonist might reduce levels of brain activity [166, 167]. (*B*) Hypothetically speaking, by exogenously modulating levels of a neurotransmitter, one might be able to titrate the dynamics of brain activity and the accompanying functional connectivity to avoid potentially damaging brain states.

Besides pharmacological drivers, there is an evergrowing body of literature suggesting that temporal fluctuations in neural acitivity and functional network patterns are largely constrained by underlying networks of structural connections [177, 178] – i.e. the material projections and tracts among neurons and brain regions that collectively comprise a connectome [179]. Over long periods of time, functional network topology largely recapitulates these structural links [180], but over shorter intervals can more freely decouple from this structure [181, 182], possibly in order to efficiently meet ongoing cognitive demands [183]. Even over short durations, functional networks maintain close structural support such that the most stable functional connections (i.e. those that are least variable over time) are among those with corresponding direct structural links [184, 185]. Multimodal and freely available datasets, such as the Human Connectome Project [186], Nathan Kline Institute,Rockland sample [187], and the Philadelphia Neurodevelopmental Cohort [188], all of which acquire both diffusion-weighted and functional MRI for massive cohorts, make it increasingly possible to further investigate the role of structure in shaping temporal fluctuations in functional networks.

## FUTURE DIRECTIONS

Prospectively, it will be important to further develop tools and models to understand the dynamic networks that support human cognition. Including additional biological realism and constraints will become increasingly important as these networks are inherently multi-layered and embedded, including spatially distributed circuits in neocortex, cortico-subcortico loops, and local networks in the basal ganglia and cerebellum. Efforts are expected to target specific computational and theoretical challenges for mathematical development including models for non-stationary network dynamics, coupled multilayer stochastic block models and dynamics atop them, and extensions of temporal non-negative matrix factorization to annotated graphs. These efforts offer promise in not only providing descriptive statistics to characterize cognitive processes, but also to push the boundaries beyond description and into prediction and eventually fundamental theories of network development, growth, and function [15, 21].

## ACKNOWLEDGMENTS

We thank Andrew C. Murphy and John D. Medaglia for helpful comments on an earlier version of this manuscript. A.N.K., A.E.S., R.F.B., and D.S.B. would like to acknowledge support from the John D. and Catherine T. MacArthur Foundation, the Alfred P. Sloan Foundation, the National Institute of Health (1R01HD086888-01), and the National Science Foundation (BCS-1441502, CAREER PHY-1554488, BCS-1631550). The content is solely the responsibility of the authors and does not necessarily represent the official views of any of the funding agencies.

## APPENDIX

Useful tools include the following:

- For dynamic community detection tools see http://netwiki.amath.unc.edu/GenLouvain/GenLouvain.
- For statistics of dynamic modules see http://commdetect.weebly.com/.
- For non-negative matrix factorization for dynamic graph models see https://github.com/akhambhati/nonnegfac.
- For statistics on dynamic graph models see https://github.com/asizemore/Dynamic-Graph-Metrics.

